# Network-based pathogenicity prediction for variants of uncertain significance

**DOI:** 10.1101/2021.07.15.452566

**Authors:** Mayumi Kamada, Atsuko Takagi, Ryosuke Kojima, Yoshihisa Tanaka, Masahiko Nakatsui, Noriko Tanabe, Makoto Hirata, Teruhiko Yoshida, Yasushi Okuno

## Abstract

While the number of genome sequences continues to increase, the functions of many detected gene variants remain to be identified. These variants of uncertain significance constitute a major barrier to precision medicine ^1–3^. Although many computational methods have been developed to predict the function of these variants, they all rely on individual gene features and do not consider complex molecular relationships. Here we develop PathoGN, a molecular network-based approach for predicting variant pathogenicity. PathoGN significantly outperforms existing methods using benchmark datasets. Moreover, PathoGN successfully predicts the pathogenicity of 3,994 variants of uncertain significance in the real-world database ClinVar and designates potential pathogenicity. This is the first computational method for the clinical interpretation of variants using biomolecular networks, and we anticipate our method to be broadly useful for the clinical interpretation of variants and for assigning biological function to unknown variants at the genomic scale.

## Introduction

As genome sequencing accelerates, the functional interpretation of biological phenomena hidden in the genomic code is becoming extremely important. Recently, genomic medicine has begun to be integrated in a clinical setting with genomic information used to optimize medical care to individual patients. Diagnostic and treatment decisions can be made based on patient genome analysis, literature, and database research on disease-related genes, alongside family history information and the patient’s clinical background. This process is referred to as the “ clinical interpretation” of genomic data. Since the information and criteria used for the clinical interpretation of variants vary from disease to disease, in actual practice, experts in each disease area discuss and decide the criteria for clinical significance based on literature information and actual cases in each hospital or institution. For example, in 2015, the American College of Medical Genetics and Genomics (ACMG) and the Association for Molecular Pathology (AMP) published guidelines for the clinical assessment of variants ^4^.

However, the functions of many detected variants from clinical sequence data remain to be elucidated. One of the reasons that variants cannot be interpreted is a lack of functional investigation, since it is unfeasible to validate the pathogenicity of such an enormous number of variants experimentally. As a result, most variants are assigned as being variants of “ uncertain significance” (VUS), which means there was no sufficient evidence to judge a relationship between the variant and disease. This is a crucial bottleneck promoting genomic medicine. Therefore, computational methods for predicting the functionality of variants are widely used ^5^, and ACMG-AMP guidelines recommend computational prediction as supporting information for variant interpretation.

A myriad of computational methods has been developed to predict deleterious non-synonymous single nucleotide variants (nsSNVs) ^6^. However, existing methods only utilize features derived from an individual variant or individual gene within the variant. In a physiological setting, genes and their products are associated with each other, and the associations have a strong influence on biological processes and disease development ^7^. These associations can be represented as biological molecular networks, which are composed of diverse biochemical and functional relationships between genes and gene products ^8^, in the form of metabolic pathways, signaling pathways, and protein–protein interaction networks. In many cases, human disease will not be caused by individual genomic alterations but by the interaction of individual mutations on multiple other pathways and relationships. It is known that utilizing molecular networks enables the identification of novel genes and pathways associated with a particular disease phenotype ^9^. Nevertheless, there are no computational methods for clinical interpretation of variants using knowledge of biological molecular networks.

## Results

### Overview of network-based prediction

To address these challenges, in this study, we developed a pathogenicity prediction model with a graph neural network (PathoGN). While existing methods predict the function of variants by machine learning using sequence and structural information of genes and variants, PathoGN achieves highly accurate prediction of variant function (pathogenicity) by learning the structure of a large biological molecular network as background knowledge. As shown in Fig. 1a, PathoGN consists of two major parts: the part that builds a large biomolecular network model by adding variant data to biological pathways and the part that learns the biomolecular network by deep learning and predicts pathogenicity of the variant.

**Fig. 1.**
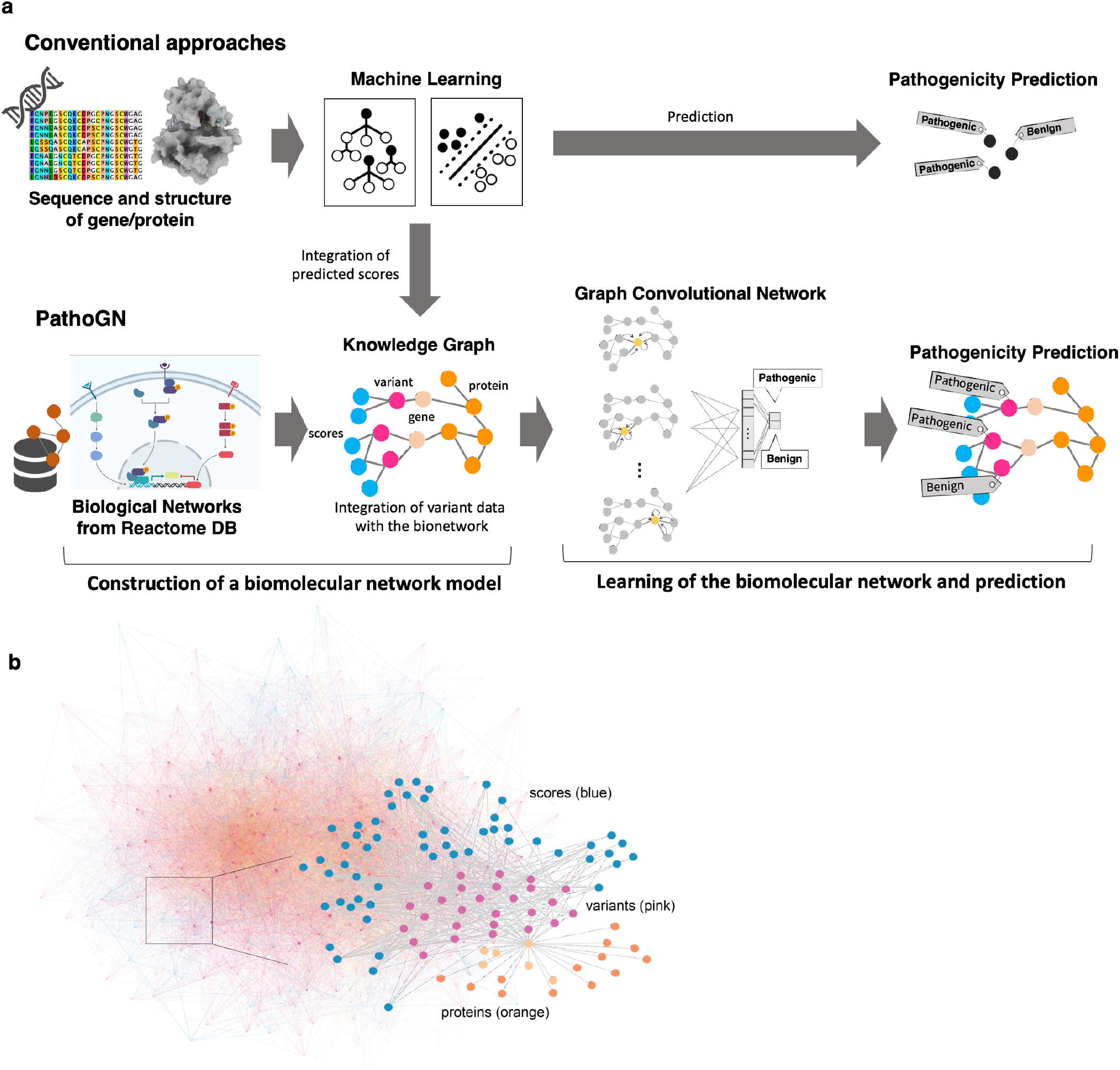
Schematic graph-based prediction method of PathoGN. (**a**) Overview of graph-based prediction. Conventional approaches use the information for individual variants such as that obtained from genome sequences. By contrast, PathoGN utilizes biological networks represented in a graph structure. Combining biological network information, disease-associated variants information, and prediction scores by conventional tools, a knowledge graph is constructed and used as an input for graph convolutional networks (GCN). Proteins, genes, variants, compounds, and prediction scores are represented by a node, and relationships between them are denoted as edges on a graph. (**b**) Knowledge graph image as input of PathoGN. The statistical detail of graph is shown in Table 1.

Table 1 shows the statistics of molecules and variants (nodes) and their connections (edges) that make up the biomolecular network (graph) used in this study. First, we constructed biomolecular networks consisting of proteins and ligands, which include protein–protein interactions, signaling pathways, and metabolic pathways from the Reactome database ^10^, a representative database of biological networks. Then, using the databases with information on effects and pathogenicity of variants, we integrated the relationships (edges) between genes and variants into the Reactome-derived network by considering the same genes and proteins to be common nodes. In addition, the pathogenicity scores predicted by the conventional method were added as nodes of the biomolecular network, resulting in the construction of a large network composed of conventional score-variant-gene-protein-ligand (Fig. 1b).

**Table 1a.** The statistics of knowledge graph for the basic biological network. The number of nodes and edges in the knowledge graph constructed from the molecular network (Reactome). Reactome instance is a Reactome record, which indicates a reaction by any of the components in the Reactome database.

| nodes |  |  | edges |  |  |
| --- | --- | --- | --- | --- | --- |
| Protein | Compound | Reactome instance | Protein - Protein | Protein - Compound | Protein-Reactome |
| 7003 | 885 | 677 | 24975 | 6805 | 2952 |

**Table 1b.** The statistics of knowledge graph for the benchmark dataset. The number of nodes and edges in the knowledge graph constructed each benchmark dataset and molecular network graph. In the node columns, fixed nodes derived from Reactome dataset are omitted. The “ total” edges include self-edges and bidirectional edges, and the “ unique” column is calculated by (total edges – the number of nodes)/2. The interaction edges (e.g. score-variant) include bidirectional edge.

|  | nodes |  |  |  |  | edges |  |  |  |  |
| --- | --- | --- | --- | --- | --- | --- | --- | --- | --- | --- |
| dataset | total | variant | gene | proteins | scores | total | unique | score-variant | gene-variant | protein-variant |
| exovar | 57192 | 8850 | 9089 | 3862 | 151 | 2646530 | 1294669 | 177000 | 17700 | 17700 |
| humvar | 98062 | 40389 | 12791 | 9488 | 154 | 3444336 | 1673137 | 807780 | 80778 | 80778 |
| predictSNP | 63286 | 16098 | 11129 | 664 | 155 | 2826576 | 1381645 | 321960 | 32196 | 32196 |
| swissvar | 62645 | 12729 | 9694 | 4830 | 152 | 2745079 | 1341217 | 254580 | 25458 | 25458 |
| varibench | 59640 | 10266 | 9521 | 4460 | 153 | 2682962 | 1311661 | 205320 | 20532 | 20532 |

Next, we used a graph convolutional neural network (GCN), a type of deep learning, to learn the biomolecular network as a knowledge graph and predict the pathogenicity of variants. GCN effectively learns the network structure by using the same concept as convolutional neural network (CNN), which has recently led to a breakthrough in image classification problems ^11–13^. While CNN features a “ convolution” process that multiplies and adds weights to the surrounding pixels of a pixel to all the pixels of an input image, GCN can learn the network structure by convoluting the relationships between the nodes of the input network along with the network structure. Our approach employs a node-focused prediction ^14^, which predicts a label corresponding to each node. Specifically, we constructed a knowledge graph in which the pathogenicity information of variants registered in the ClinVar database and VariBench datasets is attached as labels to the large molecular network constructed above. By training this graph with GCN, we can predict the pathogenicity or non-pathogenicity of variants with unknown functions, including VUS.

### Performance evaluations using a benchmark set

To investigate the effectiveness of utilizing molecular networks and associate information between variants in predicting pathogenicity, we compared the prediction performance of PathoGN and existing popular methods using a benchmark dataset.

Most of the existing methods utilize probabilistic models and machine learning algorithms with features obtained from nucleotide or amino acid sequence conservation and biochemical properties of amino acids that impact protein structure. We compared included MutationTaster ^15^, MutationAssessor ^16^, PolyPhen2^17^, Sorting Intolerant From Tolerant (SIFT) ^18^, likelihood ratio test (LRT) ^19^, and Functional Analysis Through Hidden Markov Models (FATHMM)^20^, which are commonly used in annotations with clinical interpretation. Additionally, ensemble methods combine the prediction results from multiple individual methods, improving prediction performance ^21–25^. Thus, we also compared the performance of random forest (RF) and support vector machines (SVM), which were used for ensemble learning with the same predicted scores. RF-based ensemble methods, such as REVEL^26^ and ClinPred^27^, generally show excellent performance.

Five benchmark sets by VariBench ^28^ were used for comparison. In the evaluating pathogenicity prediction methods, there was a problem with mixed training and test sets, where the variant information used to train each method was also included in the evaluation data (called type 1 circularity ^28^). VariBench provides datasets after removing duplication between datasets in each tool to solve this problem.

Fig. 2a and b show the area under the receiver operating characteristic curve (ROC AUC) and the accuracy for the six existing methods, including Mutation Taster, Mutation Assessor, PolyPhen2, SIFT, LRT, and FATHMM, and other popular algorithms, SVM and RF. For all datasets, the AUC and accuracy of PathoGN significantly outperformed all of the other methods. Notably, PathoGN also showed high accuracy in the SwissVar dataset (Fig. 2a), which contains many unpredictable variants. The difficulty of predicting the SwissVar dataset is thought to be caused by the low ratio of deleterious variants to neutral variants and the mixture of both interpretations in the same gene. PathoGN demonstrated robust accuracies even for unbalanced sets. (See Extended data Table 1 and 2 for the details accuracies and Extended data Fig. 1 for the ROC curves of all datasets)

**Fig. 2.**
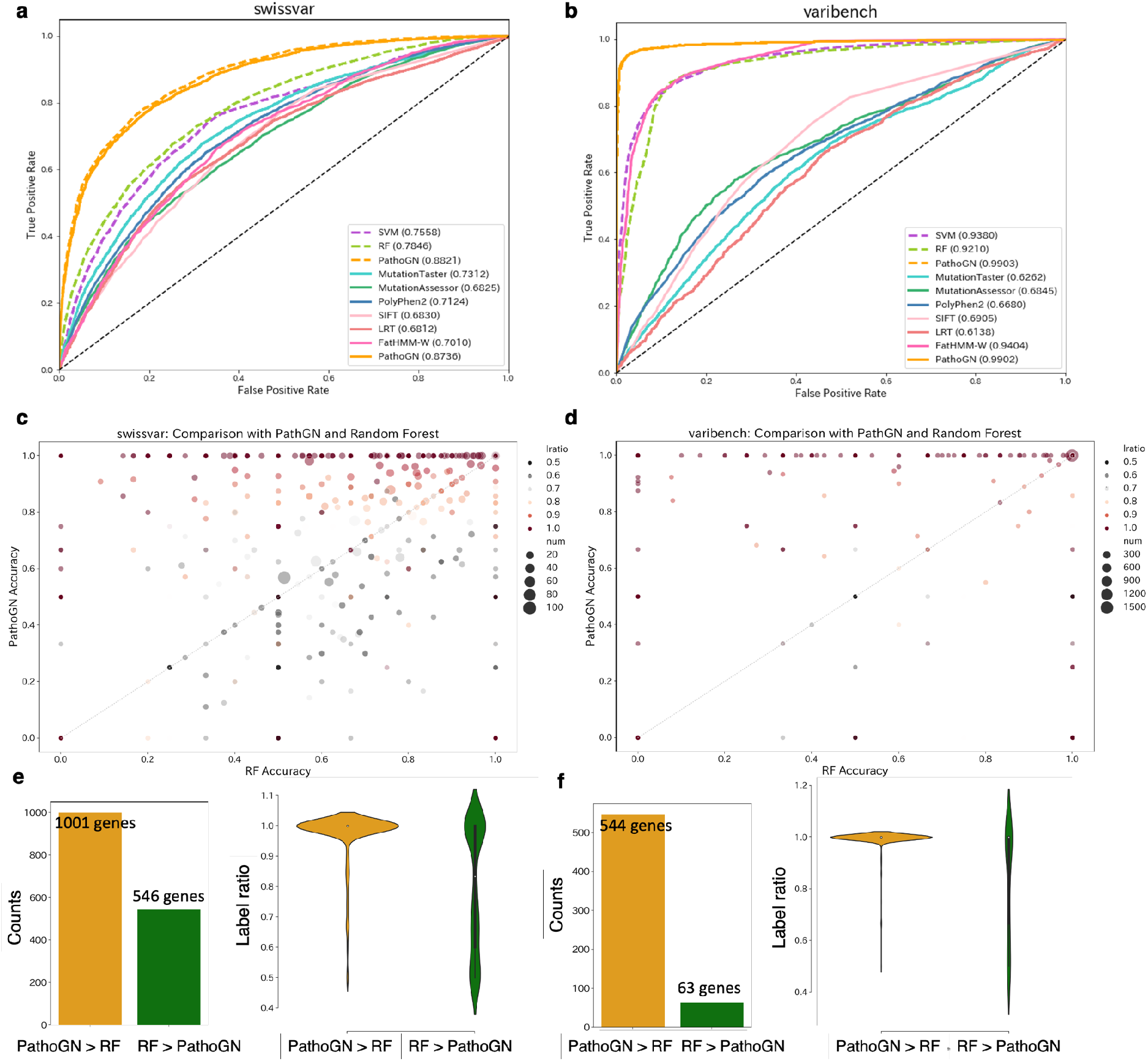
Performance evaluation. (**a, b**) ROC curves for our method (PathoGN), other non-ensemble methods, and two popular algorithms (SVM and RF) on SwissVar dataset and Varibench dataset. Solid line indicates ROC curves for PathoGN and other non-ensemble methods. For this plot, only variants for which predicted scores by all comparator tools could be obtained were used in the evaluation. Dashed line indicates ROC curve for PathoGN and other algorithms using all variants in the dataset. (**c, d**) Accuracy comparison with our approach (PathoGN) and Random Forest (RF) on SwissVar dataset and Varibench dataset. Each point represents an aggregated accuracy (mean accuracy) for each gene. The dot size represents the number of variants present in each gene in the dataset; the larger dot, the more variants on the gene are included in the dataset. The color indicates the bias of labels (label ratio) in a gene. For example, the red color indicates that all variants present in the gene have the same labels. (**e, f**) Gene counts and the label bias for PathoGN and RF. Gene were classified according to which method was more accurate (high accuracy value), and the number of genes and the bias of the labels were compared. Bar charts show the number of genes, violinplots indicate the label ratio for each gene. Orange is for genes for which prediction by PathoGN was more accurate, and green is for genes for which prediction by RF was more accurate.

In comparison with existing methods and typical algorithms, RF showed the second-best accuracy after PathoGN. Therefore, we examined the difference in prediction results between RF and PathoGN, focusing on genes. The average accuracies for all genes were calculated based on the prediction results for variants of each gene. In Fig. 2c, the comparison between the accuracies of PathoGN and those of RF in the SwissVar dataset is shown. Genes located in the upper left corner in Fig. 2c, which are more accurately predicted by PathoGN, have a higher pathogenic rate of 0 or 1, indicating that they have more similar annotations (benign/pathogenic labels) within the gene. (The comparison between PathoGN and RF for other benchmark datasets are shown in Extended data Fig. 2). Many genes were predicted with almost the same accuracies in both PathoGN and RF, but some genes were confirmed to be more accurately predicted by PathoGN (Fig 2e and f). These genes tend to have more than one variant in the same gene and have similar annotations (pathogenic/benign labels) (See also Extended data Fig. 3). Since our method enables us to use associated information between variants using the knowledge graph, these variants could be accurately predicted.

### Application to ClinVar data

To confirm the applicability of PathoGN using clinically relevant data, we evaluated the performance of PathoGN in predicting the pathogenicity of VUS with the ClinVar dataset. ClinVar is a database of genetic mutation information and its association with diseases and is widely used for interpretation in clinical practice worldwide ^29^. ClinVar updates the clinical significance of each variant when new evidence for interpretation is obtained, providing a dataset that has improved over time. Here, we performed pathogenicity prediction using past datasets of ClinVar and assessed whether these predictions were consistent with the latest annotations.

In the 2020 ClinVar dataset, 908 variants were annotated as pathogenic or benign, but were annotated as belonging to other categories (such as likely pathogenic, likely benign) in the 2019 ClinVar dataset (Fig. 3a). We refer to these variants as “ conflict variants.” We performed predictions against the conflict variants to assess how well PathoGN correctly predicts the 2020 labels. PathoGN was trained using 14817 missense variants (pathogenic: 10654, benign: 4163) in the 2019 ClinVar. Then, we used this model to make predictions for conflict variants and verified the model’s accuracy by using the labels of the 2020 dataset as the correct answer set. As a result, PathoGN successfully predicted clinical significance in the 2020 ClinVar dataset with a high predictive performance of accuracy 0.8700 and AUC 0.9410. PathoGN also showed excellent performance with the other past versions of ClinVar (2017, 2018) (Table 2).

**Figure 3.**
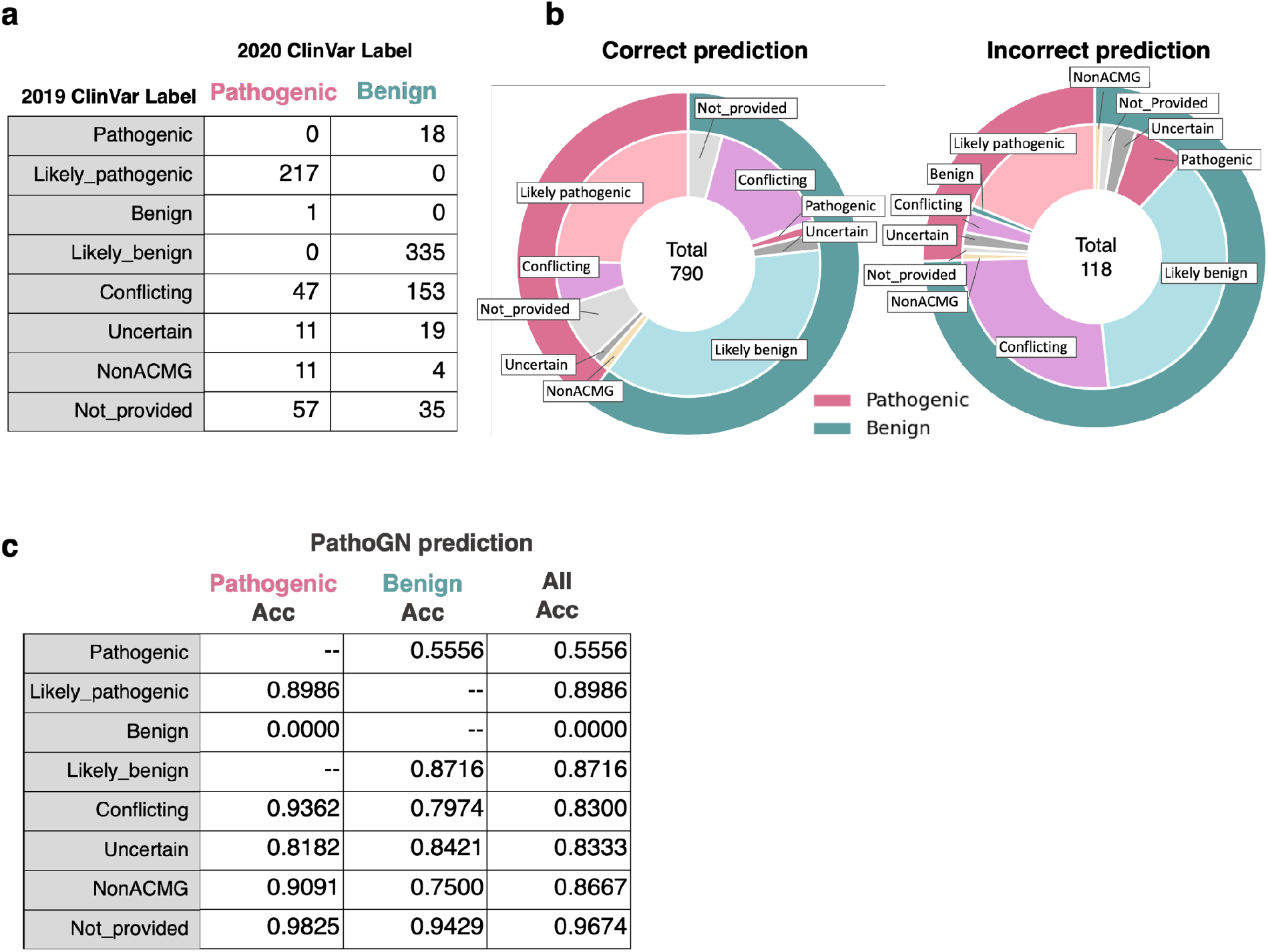
Details of predictions made with the ClinVar dataset. (**a**) Details of labels in the ClinVar dataset. Each row indicates variant’s labels in the 2019 ClinVar dataset, and the columns indicate labels assigned to the variants in the 2020 ClinVar dataset. (**b**) Each pie chart shows the category of variants. The left pie chart shows distributions of variants predicted correctly, the right pie chart shows those variants that were predicted incorrectly. The outer pie indicates the labels in the 2020 ClinVar dataset, and the inner pie shows the labels in the 2019 ClinVar dataset. (**c**) Accuracy for each significance category of variants. Pathogenic/Benign Acc column indicates prediction accuracy for the variants annotated with pathogenic/benign in 2020 ClinVar, and All Acc columns total prediction accuracy for the variants in each category.

**Table 2.** Details of the ClinVar dataset and performance results. Training set includes variants annotated as Pathogenic or Benign in ClinVar. For the “ Infer” set, variants have conflicting clinical significance among 2019, 2018, 2017, and 2020. The clinical significance in 2020 was assumed to be the correct label, and the accuracy and AUC were calculated accordingly.

|  | Training set |  | Infer set |  | Accuracy | AUC |
| --- | --- | --- | --- | --- | --- | --- |
|  | pathogenic | benign | pathogenic | benign |  |  |
| 2019 | 10654 | 4163 | 344 | 564 | 0.8700 | 0.9410 |
| 2018 | 11041 | 3893 | 409 | 727 | 0.8636 | 0.9365 |
| 2017 | 10490 | 3555 | 555 | 836 | 0.8742 | 0.9468 |

In addition, we investigated the significance category of variants for the prediction results in the 2019 ClinVar dataset (Fig. 3b and c). In all clinical significance, PathoGN shows high predictive accuracy. No correlation between label bias or the number of variants within the same gene and accuracy was observed (Extended data Fig. 4a and b). On the other hand, the accuracy tended to be lower in categories with fewer variants, such as “ Pathogenic” and “ NonACMG” than in other categories. PathoGN also enabled us to make predictions for variants with “ Not provided”. This type of variants tends to comprise those with multiple reported same significance within the same gene (Extended data Fig. 4c). This indicates that PathoGN has the ability to utilizing neighboring information from the molecular network, such as association information between variants to make these predictions.

### Predictions of VUS in the current ClinVar dataset

Next, we looked to identify the clinical significance of variants for which there is currently no known disease association. To do this, we predicted the pathogenicity for all variants that were not annotated as being either pathogenic or benign in the 2020 ClinVar dataset. The model was trained using the labeled data (pathogenic=10,877, benign=7504) and then used to make predictions for the 12,520 unlabeled variants. The details of the variants and prediction results are shown in Fig. 4a. The distribution of predicted pathogenicity probabilities for each significance category is shown in Fig. 4b. The predicted probability of being benign is calculated as one minus the pathogenicity probability. Variants annotated as likely benign in ClinVar tend to have a higher probability of being benign, whereas variants annotated as likely pathogenic tend to have a higher probability of pathogenicity. As a result, 2802 and 1192 variants with uncertain significance were predicted as pathogenic and benign, respectively. The cut-off value for the label was determined by Youden index, which is the most popular approach for determination of cut-point ^30^, and 0.3902 was used. Among them, 1570 and 593 mutations were extracted as the top 10% predicted probability of pathogenic and benign, respectively. Details of these variants and the predicted pathogenicity scores are available in GitHub and MGeND.

**Fig. 4.**
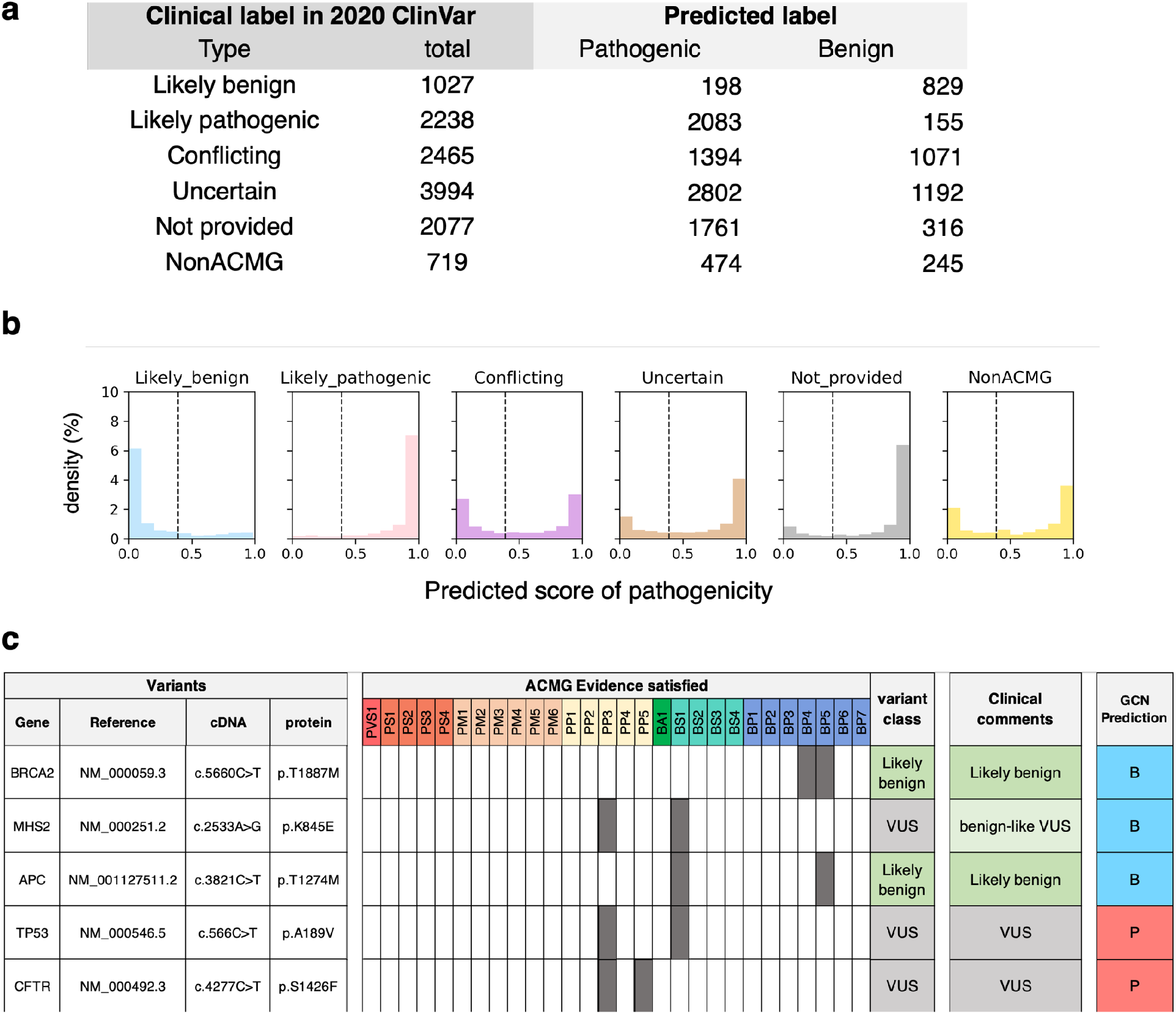
Validating predictions using ClinVar dataset. (**a**) Details of the 2020 ClinVar dataset. The “ Total” column represents the number of variants included in the 2020 ClinVar dataset and used in the prediction. Variants for which a score was available from Varibench were used in the prediction. The “ Predicted label” columns show the number of labels predicted by PathoGN. (**b**) For each significance label, the distribution of the predicted pathogenicity probability is shown. The vertical axis shows the frequency percentage normalized by the number of variants with each label, and the horizontal axis shows the predicted pathogenicity probability. Here, the predicted probability of Benign is equal to the pathogenicity probability minus one. Dash line indicates cut-off value for the label (Youden index = 0.3902) (**c**) Details of the variants reviewed with ACMG guideline. All variants are annotated as “ Uncertain significance” in 2020 ClinVar dataset.

### Validation of prediction results in the 2020 ClinVar dataset and ACMG classification

In order to evaluate the prediction results for a clinical assessment of VUS (the unlabeled variants), we performed a validation by biocurators assuming a clinical interpretation by an expert panel. Of the predicted unlabeled variants, five variants for which case information was available from the National Center for Cancer in Japan (NCC) were reviewed for classification according to ACMG guidelines. The clinical genetics professionals in NCC reviewed these variants in conjunction with the available clinical information. The reviewed variants and classification results are described in Fig. 4c.

The ACMG guidelines evaluate variants using multiple criteria and integrate each assessment to determine the variant’s final pathogenicity. The variant’s pathogenicity is represented by five categories: Pathogenic, Likely pathogenic, Likely benign, Benign, and Uncertain significance. These guidelines provide a more objective and accurate assessment.

### Variants predicted as “ Benign”

Three of the reviewed five variants were predicted to be “ Benign” by PathoGN. Firstly, BRCA c.5660C>T (p.T1887M) is assigned as BP4 because many prediction tools predicted it to be benign/tolerated/neutral. A BP5 was also assigned to this variant because BRCA1 p.E1214X, a pathogenic variant, was concurrently detected in the case. Therefore, although reported in ClinVar with VUS annotations, based on ACMG guidelines, this variant would be determined as Likely benign. Next, MSH2 c.2533A>G (p.K845E) is annotated as VUS for BS1 and PP3 criteria based on ACMG guidelines. Here, PP3 is predicted to be Pathogenic by multiple algorithms, and BS1 indicates that the allele frequency is higher than expected for the disease. The variant frequency in Human Gene Mutation Database (HGMD) is 4/2,416 (=0.001656) and in ToMMo is 23/9,546 (=0.0025). ToMMo is an integrative Japanese Genome Variation Database of the Japanese population, and the frequencies are obtained from 4.7k Japanese individuals23. HGMD manually collects and curates all published germline variants related to human inherited disease ^31^. These frequency values in the general Japanese population present a strong evidence, while the in silico prediction is a supporting evidence in the ACMG guidelines. Taken together, this variant may be considered to be a “benign-like VUS”. While many in silico prediction tools assign this variant to be Damaging/Pathogenic, PathoGN predicts “ Benign.”

The frequency of third variant APC c.3821C>T (p.T1274M) is 3/2,420 (=0.00124) in HGMD and 21/9,546 (=0.0022) in ToMMo, BS1 criteria was also assigned to this variant. Based on the publication search, it was reported that the pathogenic variant had been found in other gene involved in the molecular mechanism of the disease in a case with the same variant ^32^. From this evidence, BP5 was assigned to this variant. Thus, this variant is annotated as “ likely benign” based on BS1 and BP5 criteria. These reviews suggest that variants predicted as Benign by PathoGN may be variants that are possible to judge as Benign if more detailed information, such as functional analysis in the population, is obtained in the future.

### Variants predicted as “ Pathogenic”

Two variants predicted to be “ Pathogenic” were also reviewed for ACMG classification. For TP53 c.566 C>T (p.A189V), BS1 and PP3 were assigned, and the classification by the clinical geneticists at NCC is VUS based on the ACMG guidelines. However, the case with this variant is clinically suspected as Li-Fraumeni syndrome. For instance, the mother with breast cancer also had this variant. However, no other segregation data is available, and the biocurators could not assign other ACMG criteria to update the VUS status. The remaining variant, CFTR c.4277C>T (p.S1426F) was also reexamined and determined to be VUS.

## Discussion/Conclusion

Our novel pathogenicity prediction approach utilizing molecular networks outperformed existing methods, and the evaluation results indicated the usefulness of molecular network information as a knowledge graph for predicting variant pathogenicity. The predictions made by PathoGN also utilize variant information on the same gene. Thus, variants on genes that have hotspots can be predicted with higher accuracy. The validation using the ClinVar dataset succeeded in accurately predicting the latest clinical significance of variants whose significance was not decided in the past. The clinical decision of the variant’s pathogenicity is comprehensive, considering any information such as the family phenotype information. Therefore, it is not easy to validate the prediction results experimentally. However, PathoGN shows the possibility of making correct predictions for variants whose clinical significance is not clear at present and is expected to accelerate the re-evaluation of VUS.

Moreover, we confirmed the ACMG annotation for the variants that were predicted to be pathogenic or benign. For most of the variants, the guideline-based evidence could not be assigned and clarify their pathogenicity. The current ACMG classification is only designed for high penetrant variants. On the other hand, recent large-scale germline analyses have identified clinically significant variants in the populations that do not exhibit the classic hereditary tumor phenotype. These analyses revealed the need to discuss moderate risk variants, modifier genes, and oligogenic diseases ^33,34^. However, the clinical utility for these variants is still undetermined, and are most of them are annotated as VUS.

As a tool to support clinical interpretation, the ability to accurately predict the pathogenicity of high penetrance variants is a prerequisite. On the other hand, the interpretation of VUS in actual genomic medicine requires a more comprehensive judgment based on a thorough understanding of family phenotype and patient information. The data labeled in the current guidelines are not sufficient for this, and a range of additional information needs to be taken into account.

In the future, there is a need to expand the dataset and develop suitable models to achieve this while narrowing down the target diseases. PathoGN is suitable for such an expansion because PathoGN is the knowledge graph-based model. Incorporating detailed information, such as clinical data, into the knowledge graph, PathoGN can be easily expanded to deal with other information. This expansion may help in predictions for variants with unknown functions in the current situation. Additionally, at the moment, PathoGN is only applicable to non-synonymous single nucleotide variants (nsSNV). By improving the input network and knowledge graph, we intend to expand our model to predict insertion/deletion variants, variants in splicing sites, and variants outside of exonic regions.

The era in which individual genomes can be analyzed as a part of the standard health care services will soon be realized, and a vast amount of variants will be identified. We anticipate that the use of molecular networks and knowledge graphs will facilitate these interpretations.

## Methods

### Graph Convolutional Network model

Graph Convolutional Network model (GCN) is a type of neural network proposed by Tomas Kipf in 2017 ^35^. It takes a graph as input, convolves features of adjacent nodes, so it can learn the structural information of the graph. Networks such as molecular interactions can be represented by a graph, and GCN is suitable for network-based prediction. Here, we apply GCN on the pathogenicity prediction task. We consider an undirected graph with total *N* nodes and labeled *T* nodes.

In the pre-processing step, we construct a binary, symmetric adjacency matrix *A* ∈ ℝ^*N*×*N*^ by setting entries *A*_*i j*_ = 1, if relationships are present between nodes (i.e., proteins, genes, variants, compounds, and prediction scores) *i* and *j. Ã* = *A* + *I*_*N*_ is the adjacency matrix with added self-connections, where *I*_*N*_ is the identity matrix.

In the Embedding layer, node indices are represented by a fixed-length feature vector *x* and updated by learning. Then the values of each element are clipped to [0, 1.5]. *X* ∈ ℝ^*N*×*C*^ is a matrix of node feature vectors with dimension *C*.

In the Graph Convolution layer, node features are updated according to the following propagation rule:

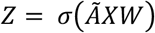

Here, *Z* is the convolved node feature matrix, *W* ∈ ℝ^*C*×*F*^ is a weight matrix, *σ*(·) denotes an activation function and 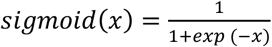 is used.

In the Dense layer, Z is projected onto binary classification probability using a fully connected layer and 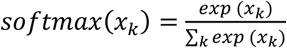.

The cross-entropy loss used as the objective function is defined as follows:

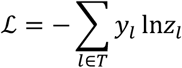

where label and classification probability are denoted by *y* and *z*, respectively.

We employed *C* = 128 and *F* = 64. We initialize weights using He Normal initialization ^36^. We train models every 1000 labeled nodes for a maximum of 100 epochs using Adam ^37^ with early stopping. The learning rate and window size of early stopping are 0.001 and 3 for the benchmark datasets, 0.01 and 5 for the ClinVar datasets. To implement this model, Tensorflow ^38^ was used.

### Benchmark datasets

We evaluated PathoGN using five datasets constructed in the VariBench benchmark set ^36^. The benchmark set consists of five datasets, *VariBench, HumVar, PredictSNP, SwissVar*, and *ExoVar*. The definition of variants as being pathogenic (damaging) or benign (neutral) vary between each dataset. The *VariBench* dataset collects nsSNV that affect protein tolerance as positive controls and obtains negative controls from dbSNP. *HumVar* dataset contains disease-causing mutations from UniProtKB database as positive controls and uses nsSNPs with MAF > 1% as negative controls. The *PredictSNP* dataset is constructed from disease-causing/deleterious variants with neutral variants from various databases such as SwissProt and HGMD. The *SwissVar* dataset collects positive controls that were found in patients or have a reported disease association from the literature, with negative controls collected from variants without any reports of disease association. In the *ExoVar* dataset, positive controls are variants related to human Mendelian disease annotated in UniProt database, and negative controls are collected using nsSNVs in the 1000 Genomes Project.

Wholly or partly *HumVar* and *ExoVar* datasets are potentially used to train MutationTester2, MutationAssessor, PolyPhen2, and FatHMM-W. Contrarily, *VariBench, PredictSNP*, and *SwissVar* datasets are provided after filtering variants used in the training phase of the prediction tools. An unavoidable exception for MutationTester2 is mentioned in ^39^. These datasets are obtained from the VariBench website (http://structure.bmc.lu.se/VariBench/GrimmDatasets.php).

### Construction of graph representation to predict pathogenicity

We defined the graph representation for ensemble prediction using protein–protein interactions obtained from Reactome ^40^ (https://reactome.org/download/current/interactors/reactome.homo_sapiens.interactions.tab-delimited.txt; Download 2021 July). The molecular network consists of 7,003 proteins, 885 compounds, 677 reactome instances, and 35,340 unique interactions. The graph consists of nodes representing each variant, gene, and protein. Each node initially has a random value. The variant nodes have labels to indicate whether they are pathogenic or benign, and prediction score nodes are predicted by MutationTaster, MutationAssesor, PolyPhen2, CADD, SIFT, LRT FatHMM-U, FatHMM-W, GERP++, and PhyloP. The prediction scores from these tools were obtained from the VariBench website, which was the same as for the benchmark datasets. We excluded Condel, Condel+, Logit, Logit+ because they include PP2, MutationAssessor, and SIFT scores. Before merging into the graph, the prediction scores were normalized to the range −8 to 8, and missing values were replaced by nodes representing a missing value.

### Validation with benchmark dataset

We constructed graphs with each dataset and trained them. For performance evaluation, prediction scores from Mutation Taster, Mutation Assessor, SIFT, PolyPhen2, LRT, and FATHMM-W in the benchmark set were used as comparators in this study. Mutation Taster ^41^ predicted the disease-causing potential of each variant with a naïve Bayes classifier that incorporated evolutionary conservation and splice-site changes. MutationAssessor ^42^ evaluated the functional impact of missense variants based on the evolutionary conservation of amino acids in protein sequences. The likelihood ratio test (LRT) ^43^ utilized a comparative genomics dataset of multiple vertebrate species to evaluate conserved amino acid positions and predicted the effect of variants. SIFT ^44^ evaluated whether variants were deleterious on sequence homology and position-specific scoring matrices with Dirichlet priors. PolyPhen2 ^45^ utilized sequence conservation and also protein information such as B-Factor and solvent accessibility and employs the naive Bayes classifier algorithm. FATHMM ^46^ incorporated hidden Markov models and sequence conservation of amino acids. FATHMM-W is an improved version that incorporated pathogenicity weights derived from disease association.

FATHMM-W was shown to have the best performance in benchmark testing by Grimm et al.^39^. Only variants for which predicted scores could be obtained by all comparator tools were used in the evaluation.

To investigate algorithm superiority, we compared performance with RF and SVM. Random forest is often employed for ensemble methods and shows good performance. For example, REVEL incorporates RF and 18 individual pathogenicity predictions and outperforms other ensemble methods. We implemented prediction models based on RF and SVM algorithms with scikit-learn. Hyperparameters of both models were selected by grid-search using 3-fold cross validation. For RF and SVM, the normalized prediction scores were also used as input features and missing values were completed by using the mean. The labels for each variant were assigned as being pathogenic or benign using cut-offs determined from the Youden index ^47^ of the predicted scores or probabilities. The Youden index is the point with the maximum value calculated by (sensitivity + specificity −1). The cut-off values for the validation are shown in Extended data Table 3.

### Validation with ClinVar dataset

The ClinVar dataset was downloaded in variant call format (VCF) from the web site of the National Center for Biotechnology Information (NCBI) ClinVar. For evaluation, a dataset downloaded on January 8^th^, 2019 was used. Missense single nucleotide variants that were annotated as pathogenic and benign were used for training, and we estimated labels for variants with conflicting significance from a dataset downloaded on February 10^th^, 2020 (2020 dataset). As some variants in ClinVar had multiple significances, we categorized variants using the following terms: Benign, Likely_benign, Likely_pathogenic, Pathogenic, Conflicting, Uncertain, NotProvided, NonACMG. Details of these categories are shown in Extended data Table 4. For each variant, we assigned the prediction scores obtained from the VariBench dataset, and variants whose scores are not listed in Varibench dataset were eliminated. The graph data was constructed using the prediction scores and Reactome dataset with the same workflow as above. For training, 14,817 variants were used, and for prediction, 908 variants were targeted. For the 2017 and 2018 datasets, the same process was performed. The datasets for 2017 and 2018 used the following versions: clinvar_20170104 and clinvar_20180401, respectively.

We evaluated our model with mean accuracy and mean AUC by 5-fold cross validation. The accuracies of each model are shown in Extended data Table 5. The model with the best performance among folds was used to estimate labels with conflicting variants. The details of conflicting variants in each dataset are shown in Extended data Table 6. The labels for each conflicting variant were decided based on the probability of pathogenicity and compared with the significance in the 2020 dataset.

### Prediction for variants with no clearly defined significance in ClinVar and manual validation

Predictions were also conducted for variants that were not annotated as being either pathogenic or benign in the 2020 dataset. The model was trained with labeled data from the 2020 dataset and then used to predict labels for the unlabeled variants. The cut-off value for prediction was determined using the Youden index ^47^. For this prediction, we employed 0.3902 as the cut-off for predictions.

Of these, variants annotated as “ Uncertain significance” in 2020 ClinVar and with more than 0.9 predicted probability of pathogenic or benign were extracted. Five variants were selected for which detailed case information could be analyzed in the National Center for Cancer in Japan (NCC). The clinical genetics professionals at NCC classified the variant pathogenicity based on the ACMG guidelines ^48^.

The ACMG guideline defines 28 criteria to assign an evidence code type for a variant. Each evidence code type is represented by benign (B) or pathogenic (P), and a level of evidence strength: stand-alone (A), very strong (VS), strong (S), moderate (M), or supporting (P). Here, multiple criteria may be applicable for a single variant. The overall combination of criteria is then used to determine if the variant is pathogenic.

## Data availability

All data are available in the main text. The knowledge graph used for PathoGN was constructed from public databases; Reactome, VariBench, ClinVar. The predicted results for all benchmark datasets and the pathogenicity scores for ClinVar 2020 dataset predicted by PathoGN are available on GitHub (https://github.com/clinfo/PathoGN.git). The pathogenicity scores for ClinVar 2020 dataset are also available in MGeND (https://mgend.med.kyoto-u.ac.jp/).

## Code availability

The open-source Python code to run the demo with PathoGN is available on GitHub (https://github.com/clinfo/PathoGN.git).

## Acknowledgments

This work was supported by the Japan Agency for Medical Research and Development (AMED) under grant number JP20kk0205013h0005. This work was supported by MEXT as “ Program for Promoting Researches on the Supercomputer Fugaku” (Application of Molecular Dynamics Simulation to Precision Medicine Using Big Data Integration System for Drug Discovery).

## Author contributions

M.K., R.K. and Y.O. conceived the ideas; R.K., M.K. and A.T. performed the prediction; M.K., A.T., Y.T., M.N., T.Y., M.H. and N.T. analyzed the experiment results. All authors contributed to the preparation of the manuscript.

