## Supplemental Figures and Tables for "Network-based pathogenicity prediction for variants of uncertain significance"

**Extended Data Fig. 1.**
**The ROC curves for our method (PathoGN), other non-ensemble methods, and other popular algorithms (SVM and RF) on all benchmark datasets.**

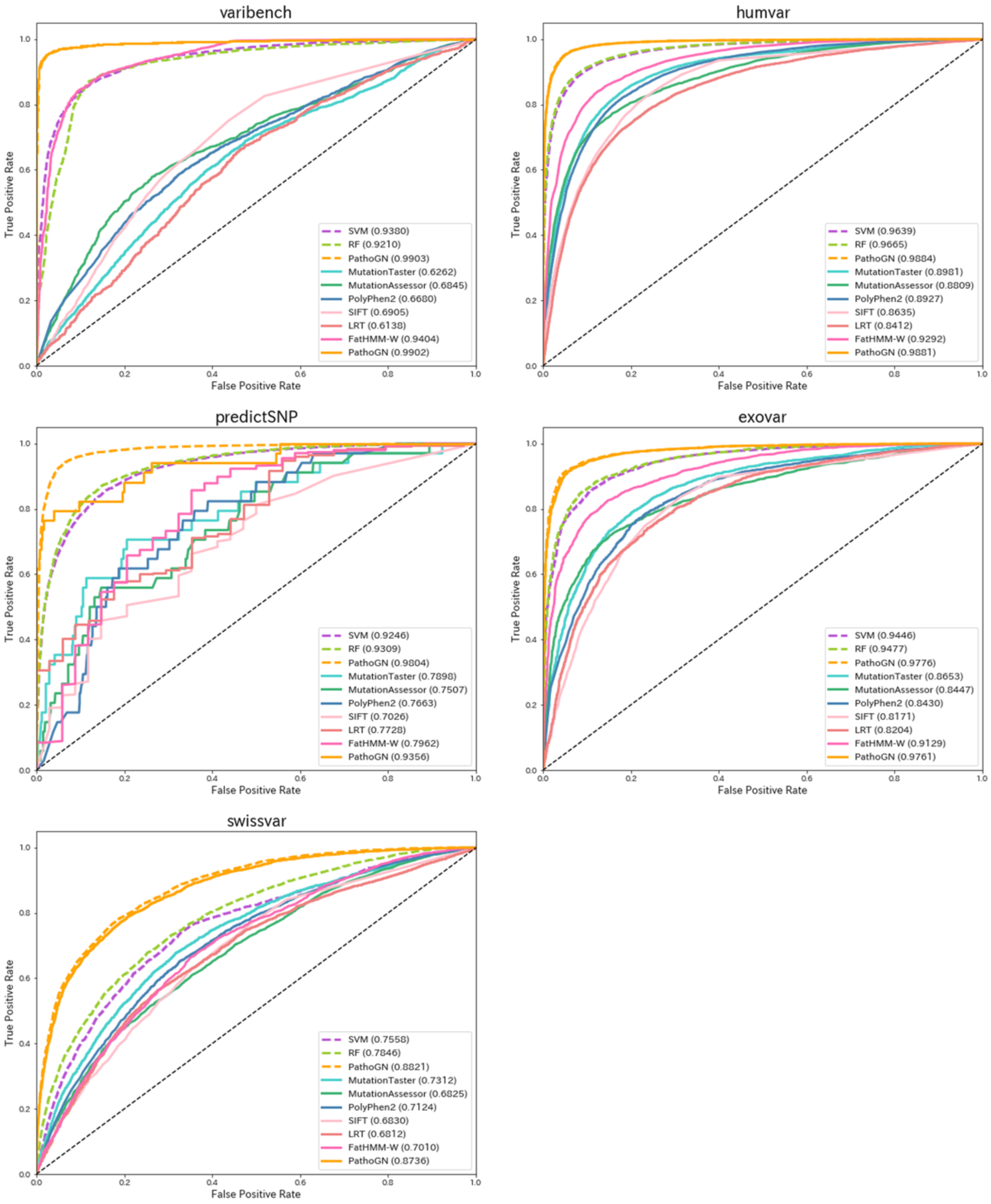


Solid line indicates ROC curves for PathoGN and other non-ensemble methods. Dashed line indicates ROC curve for PathoGN and other algorithms using all variants in the dataset.

**Extended Data Fig. 2.**
**Accuracy comparison with our approach (PathoGN) and Random Forest (RF)**

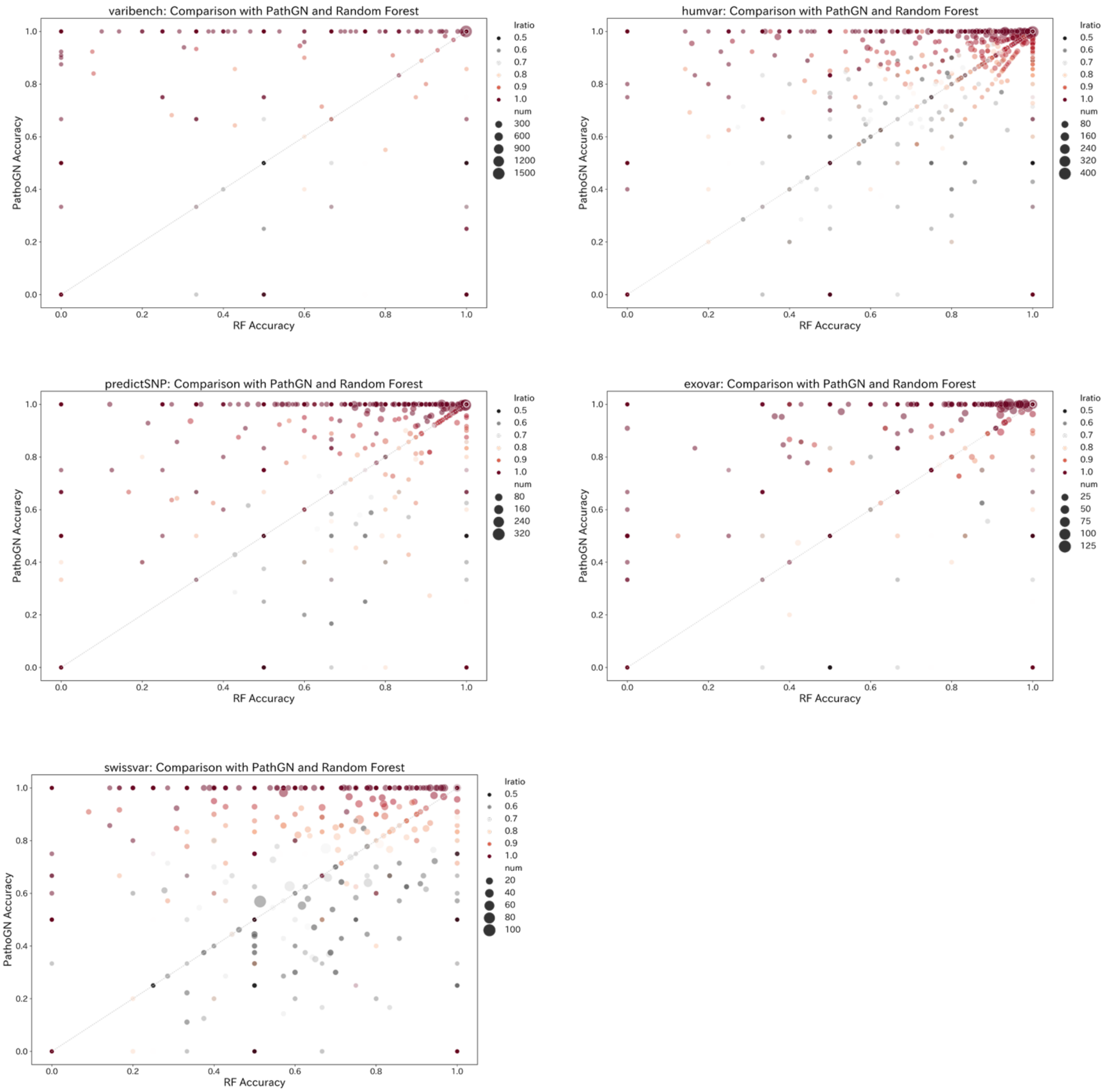


Each point represents an aggregated accuracy (mean accuracy) for each gene. The dot size represents the number of variants present in each gene in the dataset; the larger dot, the more variants on the gene are included in the dataset. The color indicates the bias of labels (label ratio) in a gene. For example, the red color indicates that all variants present in the gene have the same labels.

**Extended Data Fig. 3.**
**Violin plots of within-gene label bias for each prediction accuracy range.**

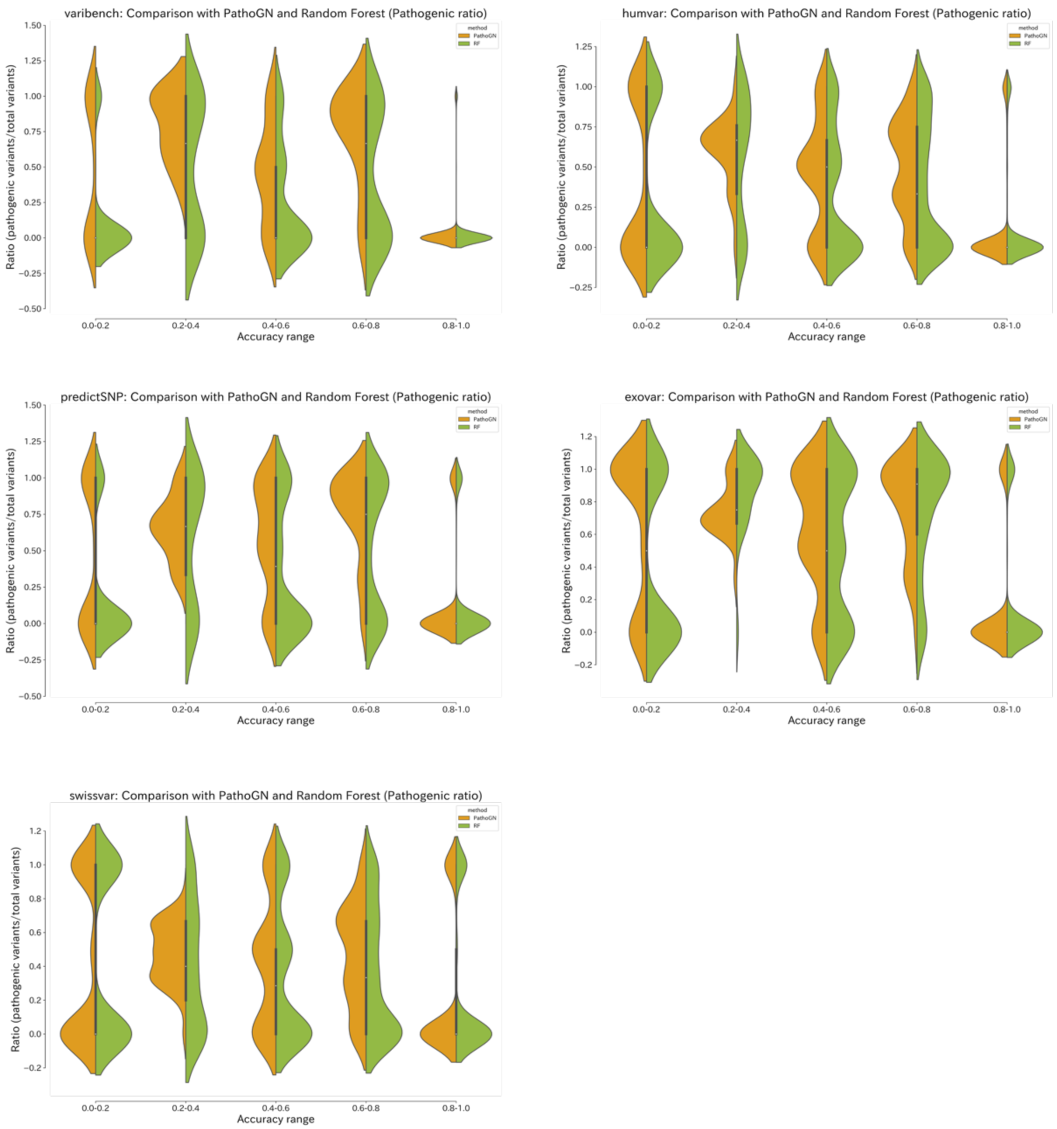


The ratio of pathogenic labels within the same gene is compared for each prediction accuracy of 0.2 increments. Orange (left) and green (right) plots are based on results from PathoGN and Random Forest (RF) predictions, respectively.

**Extended Data Fig. 4.**
**The prediction results on the 2019 ClinVar dataset**

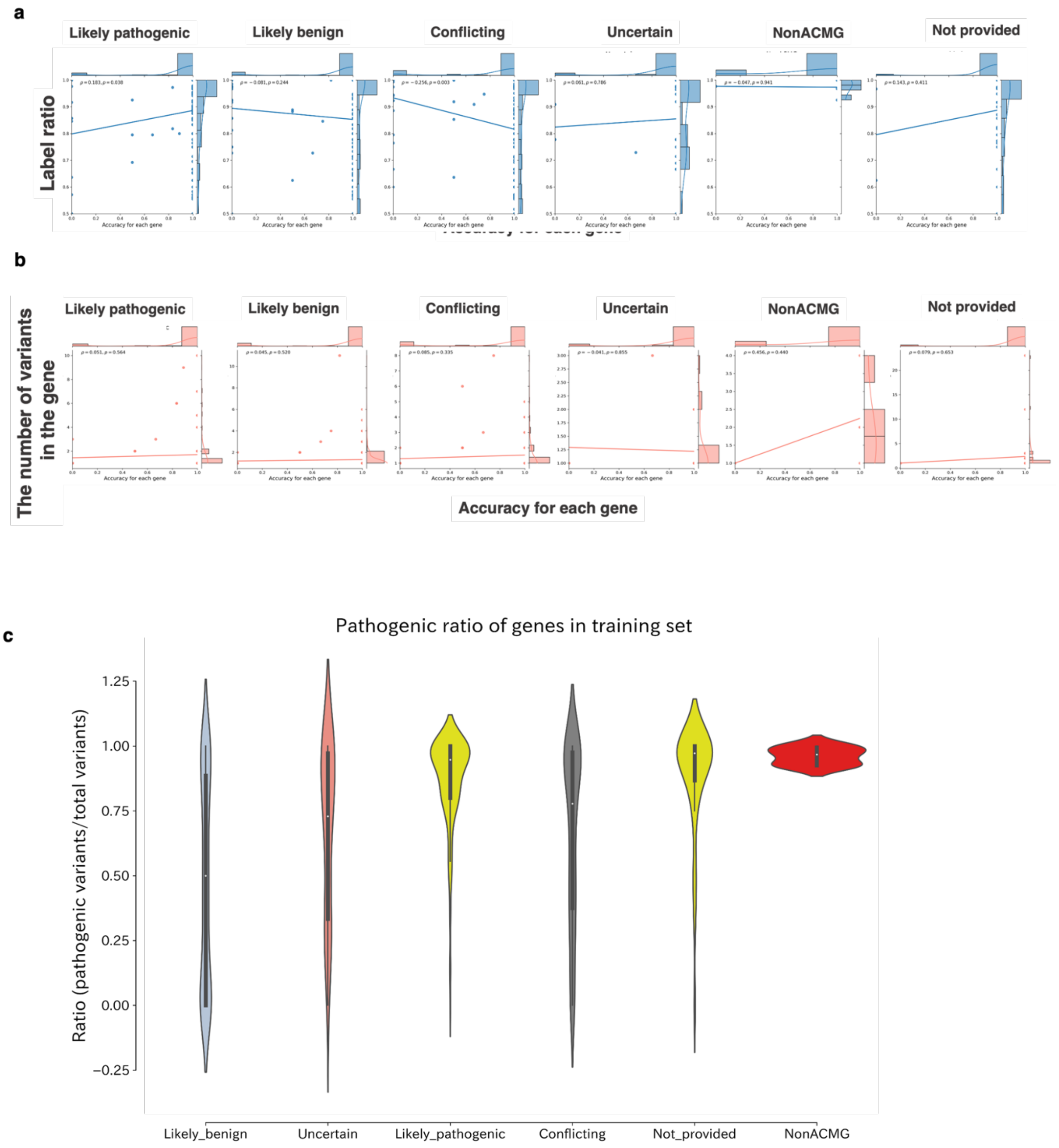


**a.** The comparison on accuracy and label bias for each clinical significance. **b.** The comparison on accuracy and the number of variants in the gene for each clinical significance. **c.** Pathogenic ratio of genes in the training dataset for each significance in the 2019 ClinVar dataset. Each violin plot shows distribution of pathogenic ratio.

**Extended Data Table 1a.**
**The details of each dataset for the comparison of accuracy with the existing methods.**

| dataset | total | positive | negative |
|---|---|---|---|
| varibench | 7994 | 3753 | 4241 |
| humvar | 31778 | 17148 | 14630 |
| predictSNP | 685 | 34 | 651 |
| exovar | 6871 | 4267 | 2604 |
| swissvar | 9451 | 3642 | 5809 |

Each dataset contains variants with positive (damaging or pathogenic) label and negative (neutral or benign) label. The "total" column indicates the number of variants for each dataset. Only variants for which predicted scores from all comparator tools could be obtained were used in the evaluation.

**Extended Data Table 1b.**
**The details of full comparison of performance with the existing methods.**

**AUC**

| dataset | MutationTaster | MutationAssessor | PolyPhen2 | SIFT | LRT | FatHMM-W | PathoGN |
|---|---|---|---|---|---|---|---|
| varibench | 0.6262 | 0.6845 | 0.668 | 0.6905 | 0.6138 | 0.9404 | 0.9902 |
| humvar | 0.8981 | 0.8809 | 0.8927 | 0.8635 | 0.8412 | 0.9292 | 0.9881 |
| predictSNP | 0.7898 | 0.7507 | 0.7663 | 0.7026 | 0.7728 | 0.7962 | 0.9356 |
| exovar | 0.8653 | 0.8447 | 0.843 | 0.8171 | 0.8204 | 0.9129 | 0.9761 |
| swissvar | 0.7312 | 0.6825 | 0.7124 | 0.683 | 0.6812 | 0.701 | 0.8736 |

**Accuracy**

| dataset | MutationTaster | MutationAssessor | PolyPhen2 | SIFT | LRT | FatHMM-W | PathoGN |
|---|---|---|---|---|---|---|---|
| varibench | 0.5913 | 0.653 | 0.626 | 0.6476 | 0.5793 | 0.8694 | 0.9653 |
| humvar | 0.8819 | 0.804 | 0.8195 | 0.7933 | 0.8091 | 0.8476 | 0.9508 |
| predictSNP | 0.6263 | 0.7708 | 0.7299 | 0.7518 | 0.6818 | 0.8657 | 0.9518 |
| exovar | 0.7957 | 0.7731 | 0.7827 | 0.7607 | 0.7722 | 0.8329 | 0.9255 |
| swissvar | 0.6317 | 0.6369 | 0.6532 | 0.641 | 0.6416 | 0.6707 | 0.7905 |

**Precision**

| dataset | MutationTaster | MutationAssessor | PolyPhen2 | SIFT | LRT | FatHMM-W | PathoGN |
|---|---|---|---|---|---|---|---|
| varibench | 0.548 | 0.6333 | 0.5926 | 0.6134 | 0.5468 | 0.828 | 0.9731 |
| humvar | 0.8691 | 0.8478 | 0.8335 | 0.8229 | 0.8199 | 0.8885 | 0.9567 |
| predictSNP | 0.0978 | 0.118 | 0.1128 | 0.1047 | 0.1102 | 0.21 | 0.5094 |
| exovar | 0.7831 | 0.8412 | 0.8218 | 0.8239 | 0.8073 | 0.8957 | 0.9635 |
| swissvar | 0.514 | 0.5266 | 0.5396 | 0.529 | 0.5257 | 0.5873 | 0.7032 |

**Recall**

| dataset | MutationTaster | MutationAssessor | PolyPhen2 | SIFT | LRT | FatHMM-W | PathoGN |
|---|---|---|---|---|---|---|---|
| varibench | 0.7386 | 0.6198 | 0.6504 | 0.6744 | 0.607 | 0.911 | 0.9526 |
| humvar | 0.9195 | 0.7761 | 0.8317 | 0.7863 | 0.8281 | 0.8206 | 0.9518 |
| predictSNP | 0.7941 | 0.5588 | 0.6471 | 0.5294 | 0.7647 | 0.6176 | 0.7941 |
| exovar | 0.9281 | 0.7823 | 0.8301 | 0.7818 | 0.8317 | 0.8273 | 0.9147 |

| | | | | | | | |
|---|---|---|---|---|---|---|---|
| swissvar | 0.8125 | 0.5708 | 0.6815 | 0.6241 | 0.7166 | 0.4893 | 0.7897 |

**F1-Score**

| dataset | MutationTaster | MutationAssessor | PolyPhen2 | SIFT | LRT | FatHMM-W | PathoGN |
|---|---|---|---|---|---|---|---|
| varibench | 0.6292 | 0.6264 | 0.6202 | 0.6425 | 0.5753 | 0.8675 | 0.9627 |
| humvar | 0.8936 | 0.8104 | 0.8326 | 0.8042 | 0.824 | 0.8532 | 0.9542 |
| predictSNP | 0.1742 | 0.1949 | 0.1921 | 0.1748 | 0.1926 | 0.3134 | 0.6207 |
| exovar | 0.8494 | 0.8107 | 0.8259 | 0.8023 | 0.8193 | 0.8601 | 0.9384 |
| swissvar | 0.6296 | 0.5478 | 0.6023 | 0.5726 | 0.6065 | 0.5339 | 0.7439 |

The threshold for PathoGN was decided with Youden index, and the values are shown in Table.S3. The labels of other tools were obtained by Varibench dataset.

**Extended Data Table 2a.**
**The details of each dataset for the comparison of accuracy with other algorithms.**

| dataset | total | positive | negative |
|---|---|---|---|
| varibench | 10266 | 4309 | 5957 |
| humvar | 40389 | 21090 | 19299 |
| predictSNP | 16098 | 10000 | 6098 |
| exovar | 8850 | 5156 | 3694 |
| swissvar | 12729 | 4526 | 8203 |

Each dataset contains variants with positive (damaging or pathogenic) label and negative (neutral or benign) label. The "total" column indicates the number of variants for each dataset.

**Extended Data Table 2b.**
**The details of full comparison of performance with other algorithms.**

**AUC**

| dataset | num | SVM | RF | PathoGN |
|---|---|---|---|---|
| varibench | 10266 | 0.938 | 0.921 | 0.9903 |
| humvar | 40389 | 0.9639 | 0.9665 | 0.9884 |
| predictSNP | 16098 | 0.9246 | 0.9309 | 0.9804 |
| exovar | 8850 | 0.9446 | 0.9477 | 0.9776 |
| swissvar | 12729 | 0.7558 | 0.7846 | 0.8821 |

**Accuracy**

| dataset | num | SVM | RF | PathoGN |
|---|---|---|---|---|
| varibench | 10266 | 0.872 | 0.8761 | 0.9661 |
| humvar | 40389 | 0.9025 | 0.9069 | 0.9517 |
| predictSNP | 16098 | 0.8508 | 0.8639 | 0.94 |
| exovar | 8850 | 0.8711 | 0.8773 | 0.929 |
| swissvar | 12729 | 0.696 | 0.7228 | 0.8023 |

**Precision**

| dataset | num | SVM | RF | PathoGN |
|---|---|---|---|---|
| varibench | 10266 | 0.8349 | 0.8386 | 0.9648 |
| humvar | 40389 | 0.9244 | 0.9302 | 0.9518 |
| predictSNP | 16098 | 0.8999 | 0.9052 | 0.9585 |
| exovar | 8850 | 0.9183 | 0.9293 | 0.962 |
| swissvar | 12729 | 0.5541 | 0.5969 | 0.6985 |

**Recall**

| dataset | num | SVM | RF | PathoGN |
|---|---|---|---|---|
| varibench | 10266 | 0.8663 | 0.8728 | 0.954 |
| humvar | 40389 | 0.8858 | 0.8884 | 0.956 |
| predictSNP | 16098 | 0.8549 | 0.8722 | 0.9443 |

| | | | | |
|---|---|---|---|---|
| exovar | 8850 | 0.8547 | 0.8543 | 0.9143 |
| swissvar | 12729 | 0.7419 | 0.6783 | 0.7815 |

**F1-Score**

| dataset | num | SVM | RF | PathoGN |
|---|---|---|---|---|
| varibench | 10266 | 0.8503 | 0.8554 | 0.9594 |
| humvar | 40389 | 0.9047 | 0.9088 | 0.9539 |
| predictSNP | 16098 | 0.8768 | 0.8884 | 0.9513 |
| exovar | 8850 | 0.8854 | 0.8903 | 0.9375 |
| swissvar | 12729 | 0.6344 | 0.635 | 0.7376 |

The thresholds for PathoGN, SVM, and RF were decided with Youden index, and the values are shown in Table.S3.

**Extended Data Table 3a.**
**Thresholds by Youden Index cut-off (Comparison with other bioinformatics tools).**

| dataset | PathoGN |
|---|---|
| varibench | 0.7686 |
| humvar | 0.5395 |
| predictSNP | 0.0555 |
| exovar | 0.7399 |
| swissvar | 0.4943 |

The cut-off value of PathoGN for each benchmark dataset on the comparison with other bioinformatic tools (Table S1)

**Extended Data Table 3b.**
**Thresholds by Youden Index cut-off (Comparison with other algorithms).**

| dataset | PathoGN | SVM | RF |
|---|---|---|---|
| varibench | 0.7077 | 0.4207 | 0.5389 |
| humvar | 0.4942 | 0.583 | 0.554 |
| predictSNP | 0.6154 | 0.6117 | 0.4819 |
| exovar | 0.7217 | 0.6338 | 0.5655 |
| swissvar | 0.4984 | 0.2721 | 0.398 |

The cut-off value of PathoGN, RF, and SVM for each benchmark dataset on the comparison (Table S2).

**Extended Data Table 3c.**
**Thresholds by Youden Index cut-off (ClinVar datasets)**

| dataset | PathoGN |
|---|---|
| ClinVar 2020 | 0.3902 |
| ClinVar 2019 | 0.7043 |
| ClinVar 2018 | 0.6954 |
| ClinVar 2017 | 0.7340 |

The cut-off value of PathoGN for the validation with the ClinVar datasets.

**Extended Data Table 4.**
**Reassignment of the clinical significance label.**

| Example of combination of clinical significance | Assigned significance label |
| --- | --- |
| Benign/Likely benign<br>Benign, risk_factor | Benign |
| Likely benign<br>Likely benign, drug response, other | Likely benign |
| Likely pathogenic<br>Likely pathogenic, association | Likely pathogenic |
| Pathogenic/Likey pathogenic<br>Pathogenic, risk factor | Pathogenic |
| Conflicting interpretations of pathogenicity<br>Conflicting interpretations of pathogenicity, Affects | Conflicting |
| Uncertain significance<br>Uncertain significance, drug response | Uncertain |
| not provided | NotProvided |
| Affects<br>association, risk factor | NonACMG |

Reassignment of labels for the variants with multiple combinations of clinical significance. In the ClinVar dataset, multiple significances are often assigned to a variant. For these variants, in this study, we reassigned the category label as shown in the table.

**Extended Data Table 5.**
**Validation on the training dataset.**

| version | Acc_Mean | AUC_Mean |
|---|---|---|
| ClinVar 2020 | 0.9073 | 0.964 |
| ClinVar 2019 | 0.8955 | 0.9585 |
| ClinVar 2018 | 0.8961 | 0.9579 |
| ClinVar 2017 | 0.8914 | 0.9557 |

The evaluation of the trained model for each ClinVar dataset. The trained PathoGN models were evaluated by mean accuracy and mean area under the ROC curve (AUC) via 5-fold cross-validation in the training dataset.

**Extended Data Table 6.**
**Details of target variants (conflicting variants) in ClinVar datasets.**

| | **2019 dataset** | | **2018 dataset** | | **2017 dataset** | |
|---|---|---|---|---|---|---|
| | **2020 ClinVar label** | | **2020 ClinVar label** | | **2020 ClinVar label** | |
| Previous ClinVar Label | **Pathogenic** | **Benign** | **Pathogenic** | **Benign** | **Pathogenic** | **Benign** |
| Pathogenic | 0 | 18 | 0 | 27 | 0 | 26 |
| Likely_pathogenic | 217 | 0 | 169 | 0 | 269 | 1 |
| Benign | 1 | 0 | 1 | 0 | 1 | 0 |
| Likely_benign | 0 | 335 | 0 | 385 | 0 | 481 |
| Conflicting | 47 | 153 | 84 | 250 | 67 | 238 |
| Uncertain | 11 | 19 | 16 | 15 | 25 | 24 |
| NonACMG | 11 | 4 | 44 | 8 | 48 | 16 |
| Not_provided | 57 | 35 | 95 | 42 | 145 | 50 |

The details of conflicting variants in each ClinVar dataset. Each row indicates the variant's labels in 2019, 2018, and 2017 ClinVar dataset, and the columns indicate variant labels according to the 2020 ClinVar dataset.
